# The hippocampus and cortical memory networks have an inflection point in middle childhood

**DOI:** 10.64898/2026.08.19.745842

**Authors:** Lena J. Skalaban, J. Benjamin Hutchinson, Vishnu P. Murty

**Author notes:** Corresponding Author: Lena J. Skalaban, e.

## Abstract

Decades of developmental memory research has mainly reported linear and protracted changes in both human hippocampal function and connectivity between the hippocampus and cortex. While foundational, very few studies have interrogated the reliability of hippocampal signals across age, and how this coincides with (or diverges from) age-related changes in connectivity to broader cortical networks supporting multiple memory systems. Here, utilizing movie-watching fMRI data in children 3-12 years and adults, we assessed hippocampal response stability using an inter-subject functional correlation (ISFC) approach, and then measured functional connectivity between the hippocampus and the Posterior Medial (PM) - Anterior Temporal (AT) cortical memory networks proposed to support episodic-like (PM) and semantic-like (AT) memory respectively. Results showed that hippocampal responses are stable in the youngest children, but bifurcate in 7-year-olds –– with half the subjects correlating most highly with younger and half with older age groups. Likewise, we found that while functional connectivity *within* the AT network is stable across development, connections *between* the anterior hippocampus and this network did not reach adult levels until around 7-years. Thus, while brain networks supporting semantic memory may be in place early, interactions with the hippocampus may not develop until after middle childhood –– with an inflection point around 7-years of age.

**Significance Statement:** The development of the hippocampus, a key-region in episodic memory behaviors, extends well into adolescence. However, little is known about the stability of hippocampal responses across age and how this might impact communication with cortical brain networks supporting episodic and semantic memory. Here, we show that connectivity *within* a network that supports semantic memory matures early, but connectivity *between* this network and hippocampus does not mature until at least 7-years of age. Unlike all other ages, 7-year-olds’ hippocampal responses are not stable: half of the subjects’ responses appear immature and half mature for their age. This work identifies 7-years as a developmental inflection point in both hippocampal signaling and hippocampal-cortical memory interactions that may support multiple memory systems.

## INTRODUCTION

The human hippocampus shows rapid developmental change in early childhood protracted through adolescence (Keresztes et al., 2017; Lee et al., 2017; Plachti et al., 2023). Decades of work has shown largely linear structural (Canada et al., 2020; Gogtay et al., 2006; Lee et al., 2014) and functional (Geng et al., 2019; Ghetti & Bunge, 2012) changes in the hippocampus, particularly along its’ long-axis (Blankenship et al., 2017; DeMaster et al., 2014; Nichols et al., 2023; Vijayarajah & Schlichting, 2024). Many of these developmental shifts in hippocampal structure and function have been interpreted as reflecting the slow development of episodic memory systems over time – – suggesting a unified function for the hippocampus that does not mature until adulthood. However, we know that the hippocampus does not serve a single functional purpose (Eichenbaum et al., 1992; Poppenk et al., 2013), but rather supports diverse behaviors that emerge during different periods of development. For example, hippocampal function may support pattern completion and generalization during early childhood (Keresztes et al., 2018; Ngo et al., 2018), and then transition to more pattern separation processes later in childhood (Canada et al., 2019; Cohen et al., 2025), though exactly how and when these functions develop is complex (DeMaster & Ghetti, 2013). These developmental transitions in memory functions may be influenced by changes in hippocampal response properties, as well as changes in communication with broader cortical networks to support more than one memory system. However, little work has interrogated whether developmental changes in the functional properties of the hippocampus are concomitant with age-related changes in connectivity between the hippocampus and broader cortical memory networks.

Hippocampal function changes drastically, but gradually across the lifespan. Seminal work demonstrated that univariate activation changes most rapidly from early to middle childhood (Ghetti et al., 2010), with continued increases in response specificity to certain memory tasks across adolescence (Sastre III et al., 2016; Selmeczy et al., 2021). These changes have also shown differences along the hippocampal long-axis: the posterior hippocampus appears to be engaged earlier in development, though refinement continues throughout childhood (Callaghan et al., 2021) with anterior hippocampal activation related to memory performance not mirroring adult-levels until later childhood or adolescence (DeMaster & Ghetti, 2013; Langnes et al., 2019). Likewise, though recent multivariate work on hippocampal representations has revealed that even young children can show adult-like pattern separation (Benear et al., 2022), activation patterns in the posterior hippocampus specifically shows age-related changes in pattern differentiation (Callaghan et al., 2021), with later refinement/recruitment of the anterior hippocampus during late childhood/adolescence to support adult-like episodic memory behaviors (DeMaster et al., 2016; Xie et al., 2024).

This foundational work often uses an analysis approach that measures brain responses averaged within each age group or models linear developmental change. While valuable, this approach cannot answer questions about the reliability and synchrony of responses within and across developmental stages that may reveal subtler non-linear developmental changes. Indeed, recent theoretical frameworks (Bauer, 2015; Keresztes et al., 2018), suggest that hippocampal function may be optimized to reflect the demands of the system within a given developmental period rather than developing linearly to an adult endpoint. To investigate these questions, here, we opted for an Inter-subject Functional Connectivity (ISFC) approach, which has recently been utilized to characterize differences in the response properties of the hippocampus across age-groups (Cohen et al., 2022) and generally can speak to the reliability of functional responses in a given brain region (Nastase et al., 2019). This approach allows us to arbitrate two competing hypotheses of hippocampal development: if hippocampal function matures linearly, then we should expect less ISFC in younger groups that increases slowly over time. However, if hippocampal function is a process that reflects the idiosyncratic demands of a given age-group, we should see consistent variability within group which may change as a function of age.

Though this ISFC approach can zoom-in on the developmental profiles of hippocampal functions, it cannot interrogate how information processed in the hippocampus is communicated to the cortex to support multiple memory systems. Complementary learning systems theory in adults posits that information processed in the hippocampus is transferred to the cortex over time for longer-term storage and generalization (Kumaran et al., 2016; McClelland et al., 1995; Norman & O’Reilly, 2003). Coordinated activation of the hippocampus and these cortical networks supports future retrieval of both episodic and semantic memories (Ranganath & Ritchey, 2012; Robin & Moscovitch, 2017). During development, the hippocampus has been shown to functionally integrate with widespread cortical regions that may support semantic memory around middle childhood (Blankenship et al., 2017) which is in turn is associated with improvements in episodic memory (Riggins et al., 2016). Similarly, structural changes in the hippocampus that extend into middle childhood have been related to improvements in pattern completion/separation behaviors that may support semantic extraction (Keresztes et al., 2018). In contrast, regions supporting behaviors related to semantic memory, including the anterior temporal cortex, show relatively stable functional organization from early childhood (Dehaene-Lambertz et al., 2018; Richardson et al., 2018).

However, few studies have directly interrogated developmental changes in the hippocampus and broader cortical networks in a wide age-range to untangle how and when they start integrating with one another. We wanted to extend our approach outside of the hippocampus to look at large-scale memory networks centered on the hippocampus: namely the PMAT network (Barnett et al., 2021; Ritchey et al., 2015) that is hypothesized to subserve related by separable memory functions. The posterior medial (PM) subnetwork has been shown in adults to support recall of episodic details (Cooper et al., 2021; Cooper & Ritchey, 2019; Ritchey et al., 2015) and event segmentation during movie watching (Barnett et al., 2021). In contrast, the anterior temporal (AT) network supports broader semantic/gist-like representations (Ranganath & Ritchey, 2012) and affective memories (Cooper & Ritchey, 2019). We selected these networks because of the tension between the timing of availability of semantic-features relative to episodic features across development. The hope is that differential connectivity profiles in this PM versus AT network could further elucidate the nature of what content may be processed across the hippocampus at different developmental stages. While a recent study has begun to characterize the PMAT network in older youth 8-21 years (Xie et al., 2024), research has yet to narrow-in on patterns of change in these networks during the key transitions between early to middle childhood, and how they synchronize with age-related changes in hippocampal function.

Here, we wanted to explore hippocampal function through a lens that is focused on relative development within each age group, and how these systems shift in their functional orientation across age. To explore these processes, we re-analyzed a publicly available movie-watching fMRI data from children aged 3–12 years and adults (Richardson et al., 2018) that allowed us to explore two different functional profiles of the hippocampus within and across age groups. Ideally, we would also be able to behaviorally probe in real-time individual subjects’ memories. However, this relies on communication skills that vary widely across development and thus are inherently limited by the nature of the memory tests being administered. Movie watching data however can time-lock brain activation responses to a stimulus across and generally has more compliance in functional MRI with younger children (Vanderwal et al., 2015). Furthermore, fMRI often uses sparse, static stimuli, that may not express the full dynamic range of hippocampal function (Finn & Bandettini, 2021), whereas movie-watching data in adults has been tied to a range of relevant memory functions including novelty detection (Antony et al., 2021; Ben-Yakov & Henson, 2018), internal marking of event boundaries (Baldassano et al., 2017; Silva et al., 2019) and passive encoding (Hasson et al., 2008). Although observing these brain networks independent of behavior can give only indirect proxies of episodic and semantic memory behaviors, we propose that this approach still gives valuable insight into the building blocks of brain networks necessary for processing information in both memory systems.

Utilizing the movie-watching data in a large age-range, we conducted ISFC analyses within- and across-group patterns of hippocampal activity to assess age-related changes in the consistency of hippocampal responses. We then used seed-based functional connectivity to assess age-related changes in how the hippocampus and the PMAT network share information with one another. We hypothesized that (1) hippocampal responses to the movie would show increasing stability with age as measured by ISFC (2) connectivity within the cortical semantic network would be relatively stable across childhood, (3) connectivity between the hippocampus and this network would show linear protracted development, with particular specificity to anterior hippocampus.

## RESULTS

### Overview of Analyses

All analyses were conducted using the functional timecourses of 150 subjects who watched a short Pixar movie (“Cloudy”) in the scanner (Richardson et al., 2018). We split this sample of subjects into five age-groups: 3-4 year-olds (N = 31); 5-year-olds (N = 34); 7-year-olds (N = 23); 8-12 year-olds (N = 34) and Adults (N = 28). The age-groupings were created to best balance the number of subjects in each age-group while also representing key timepoints in the development of memory.

First, to investigate how the hippocampus processes information across development, we took an ISFC approach *within* age groups to demonstrate any change in the degree of consistency in hippocampal responses using a whole bilateral hippocampal ROI. We then followed up in the 7-year-old age-group only with an *across* age group ISFC approach that calculated which age-group individual subjects correlated with most highly (using a winner takes all approach). We did this as an exploratory analysis to try and explain why only 7-year-olds, but no other age-group, showed *negative* inter-subject synchrony with their own age-group average.

Next, to investigate how the hippocampus integrates information with a cortical semantic memory network, we used a seed-based functional connectivity approach. We first looked at the age-group averaged connectivity *within* the regions of the AT network to assess the maturity of the semantic memory network alone. We then looked at connectivity *between* the whole bilateral hippocampus and AT network to assess integration between episodic and semantic information. As a control set of regions, we also looked at connectivity within/between a cortical posterior medial network (PM) and the whole hippocampus. The PM network in adults, while also highly integrated with the hippocampus, has been shown to subserve distinct episodic memory functions from the AT network (Barnett et al., 2021). Finally, in a post-hoc follow-up, we split the hippocampus along its’ long-axis (anterior and posterior) to assess whether any developmental differences observed in connectivity from these cortical memory networks to the hippocampus showed anterior/posterior specificity.

### Hippocampal Inter-subject Correlation Within Age Group

To assess whole hippocampal synchrony by age, we first conducted an inter-subject correlation analysis using each subjects’ hippocampal timecourse within a given age group. We then ran a one-way ANOVA to determine if there were any differences between age groups’ averaged Fisher z-scored correlation values. A one-way ANOVA showed a significant difference between the age groups (*F*(4,150) = 11.31, *p* <.001). Post-hoc Turkey’s HSD’s showed that 7-year-olds’ inter-subject hippocampal correlation was on average negative (*M* = −0.06, *SD* = 0.15) and significantly lower than all other age groups both younger –– 3-4 years: (*M* = 0.18, *SD* = 0.23, *p* = .001, CI = [-0.390, - 0.072]); 5 years: (*M* = 0.32, *SD* = 0.22, *p* <. 001, CI = [-0.533, −0.220]), and older –– 8-12 years: (*M* = 0.15, *SD* = 0.15, *p* = 0.004, CI = [0.046, 0.358]); Adults: (*M* = 0.20, *SD* = 0.27, *p* = 0.004, CI = [0.088, 0.413]) (See Figure 1A).

**Figure 1.**
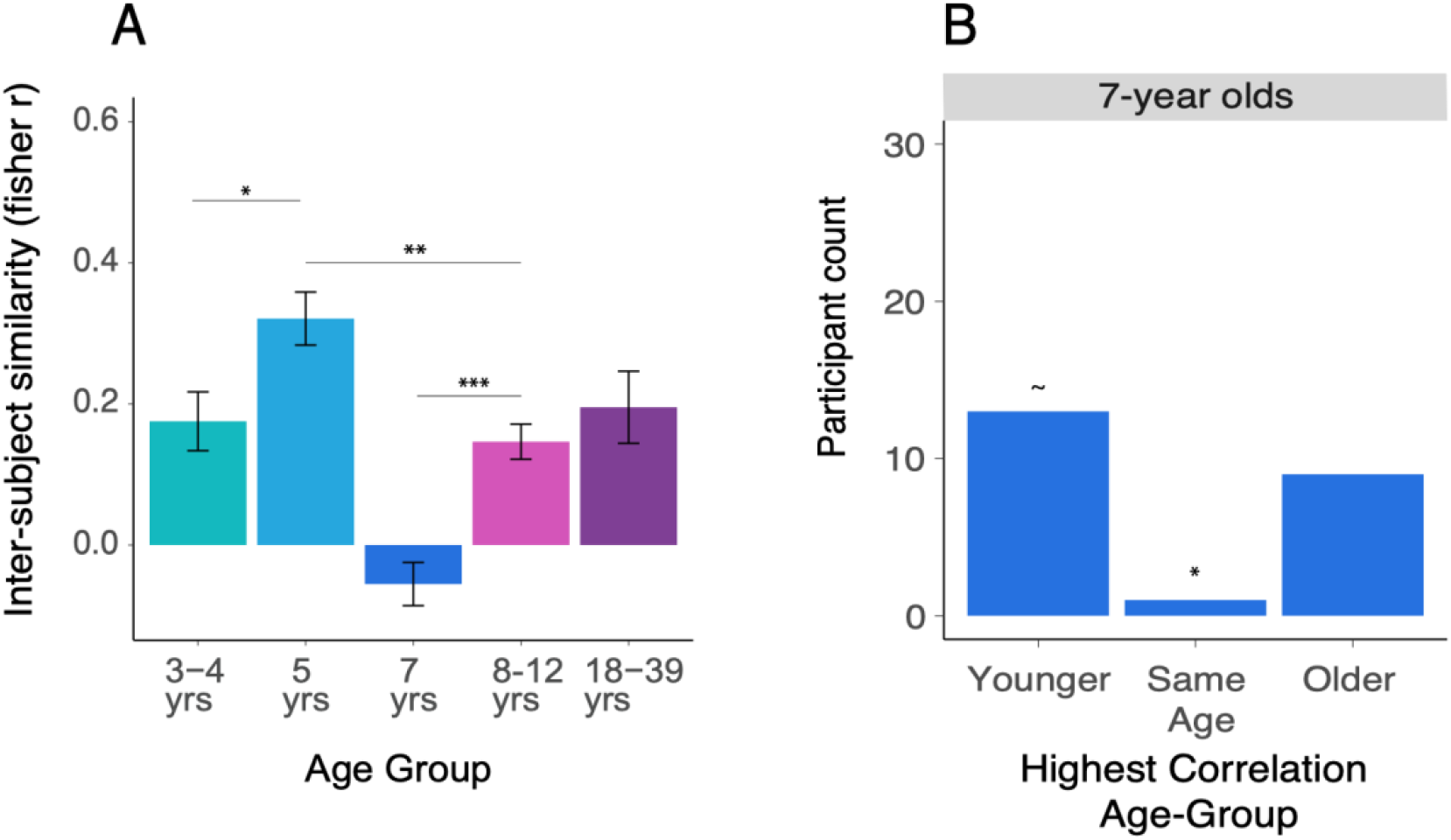
Inter-subject correlation in the whole hippocampus. A). Within age-group inter-subject correlation results. A one-way ANOVA revealed significant differences between our five age-groups. Follow-up Tukey’s HSD test revealed that there were significant differences between 7 year old age group that had a negative inter-subject fisher r value on average and all other age groups that showed positive synchrony on average. B). Histogram showing 7 year old’s highest correlations grouped based on whether those correlations were with a younger, older, or their same age group. Permutation test results revealed that there were fewer than expected 7 year olds who were most correlated with their own age group than would be expected by chance. Significance markers indicate results of simple effects tests for A and permutation tests for B (*p*< .05 *; *p*< .005 **, *p*<.0005 **).

We also found that inter-subject correlations in the hippocampus increased slightly from 3-4 years to 5 years of age (*p* = 0.045, CI = [0.002, 0.289]) and that correlations in this 5-year age-group were also higher than the 8-12 year old age group on average (*p* = 0.007, CI = [-0.315, −0.0344]). Beyond this, no other significant differences between age groups were observed –– 3-4 and 8-12 year olds: (*p* = 0.981, CI = [-0.173, 0.115]); 3-4 year olds and Adults: (*p* = 0.996, CI = [-0.131, 0.171]); 5-year-olds and Adults: (*p* = 0.134, CI = [-0.273, 0.022]) 8-12 year olds and Adults: (*p* = 0.891, CI = [-0.099, −0.196]).

In summary, while 5-year-olds’ have slightly more consistency in their hippocampal responses than other non-adult age groups, there is relatively consistent hippocampal synchrony across development –– except at 7 years of age when hippocampal responses become briefly, but markedly asynchronous.

### Hippocampal Inter-subject Correlation Between Age Groups

Given the surprising lack of synchrony in the 7-year-olds compared to all other age groups, we next conducted ISFC analyses between 7-year-old subjects and *all* age groups to try to determine if the 7-year-old participants correlated most with any particular age-group besides their own.

To do this, we adopted a winner-takes-all approach where we compared every 7-year-old subjects’ hippocampal timecourse to the average timecourse of each age-group including their own (in which case their own data was left out of this average). We labeled only the highest correlation value (winner takes all) for each subject by the according age group. We plotted a histogram summary counting the number of highest correlation values according to whether that value belonged to a younger, the same, or older age group average than that subjects’ actual age (Figure 1B). We found that the 7-year-old sample bifurcated; over half of the 7-year-olds were most highly correlated with a younger age group (13/23), and slightly less than half were most highly correlated with an older age group (9/23). However, only a single subject (1/23) had a hippocampal response most highly correlated with their own age group. While this could explain why the inter-subject correlation for 7-year-olds was negative, it is possible that the observed pattern of higher correlations with two age groups younger or older was simply due to chance, as there were twice as many age groups for them to correlate with.

### Permutation Test of Winner-Takes-All Results

To evaluate whether (1/23) was fewer 7-year-olds than would be expected by chance and as a control analysis, we conducted montecarlo permutation analysis assuming that each 7-year-old was equally as likely to be correlated with any of the other five age groups. We simulated this process for the same number of subjects each time randomly assigning which age group the simulated participant was most highly-correlated with and re-labeled as “younger”, “same age” or “older”. This produces a null distribution to which we could use to assess whether our observed pattern of results for the 7 year olds arose by chance.

The omnibus permutation test across the 3-bins: Younger, Same Age, and Older classifications was not significant (permutation test, *p* = 0.119), indicating that the overall three-category distribution did not differ reliably from chance expectations. However, because our specific question concerned whether children were likely to resemble their own age group, we also conducted a focused permutation test comparing Same-age classifications against all Different-age classifications. Indeed, in this more specified test, there were significantly fewer 7-year-olds most highly correlated than their own age group than would be expected by chance (*p* = 0.038), as well as a trend towards more 7-year-olds that were most highly correlated with younger age groups than would be expected under the null (*p* = 0.079), but no more subjects that correlated most with older age groups as would be expected under the null (*p* = 0.617).

While the overall pattern of results showing the bifurcation should be interpreted with caution, as the omnibus permutation test was not significant, likely due to the smaller sample size (n = 23) in this age-group, this simulation demonstrated that there were significantly fewer 7-year-olds that looked like their own age-group than expected by chance. This offers a potential explanation as to why 7-year-olds hippocampal responses appeared asynchronous. 7 years of age may be a hippocampal inflection point where children (except a single participant) appear to have hippocampal responses that either lag behind (are developmentally immature) or outpace (are developmentally mature) their age.

Hypothetically, this developmental bifurcation could be due to individual differences in demographic factors. The dataset also contained information on KBIT scores (age-standardized IQ), scores from a battery of Theory of Mind (ToM) tasks (% correct out of 24 questions) and gender. We tested the whether the 22 7-year-old children who most correlated with younger or older participants (leaving out the 1 subject who correlated mostly highly with their own age group) differed in terms of these demographic factors. The mean KBIT scores of the participants who looked older (*M* = 120.78) did not differ from the KBIT scores from those who looked younger (*M* = 117.15), in a Welch two-sample t-test: (*t*(22) = −0.474, *p* = 0.64). The mean ToM scores also did not differ between those who appeared younger (*M* = 0.87) and those who appeared older (*M* = 0.86) in a Welch two-sample t-test: (*t*(22) = 0.292, *p* = 0.78). Finally, there was no difference in the number of participants who were identified as male and female for the group that appeared younger (*Female* = 5, *Male* = 3), and the group who appeared older (*Female* = 6, *Male* = 8) using a Fisher’s exact test (*p* = 0.387). Thus, the differences observed between those participants whose hippocampal responses appear more or less mature may be due to other factors not measured in this dataset or simply exist independent of demographic differences.

### Connectivity with the PMAT Network

To assess how the hippocampus communicates with regions important for semantic and episodic memory respectively, we conducted a seed-based connectivity analysis using regions in the PMAT network. This network of cortical memory regions can be further split into the PM (Posterior Medial) and AT (Anterior Temporal) subnetworks that have been shown in adults to support related but separable episodic and semantic memory functions (Barnett et al., 2021) and connect to the posterior and anterior long-axis of the hippocampus respectively (Ritchey et al., 2015).

One concern with these a-priori ROI seed-based analyses might be that any age-related changes we are seeing in our regions of interest are spurious: the same age-related changes might be seen across any ROI in the brain. To address this, we first ran a control analysis looking at connectivity between left and right precentral gyrus (as a proxy for bilateral early motor cortex) as we hypothesized that there would be no difference in connectivity amongst our age groups for motor regions during a movie watching task. We ran an ANOVA assessing connectivity between left and right precentral gyrus by age-group, and indeed found that there was no significant effect of age-group (*F*(4,150) = 0.4132, *p* = 0.166). Furthermore, all age groups showed expectedly high average fisher’s r correlation values between left and right precentral gyrus (3-4 years: *M* = 1.01, *SD* = 0.57; 5 years: *M* = 0.99, *SD* = 0.51; 7 years *M* = 1.15, *SD* = 0.45; 8-12 years *M* = 1.26, *SD* = 0.54; Adults *M* = 1.15, *SD* = 0.38). The results of this control analysis reduces the likelihood that any low functional connectivity values was due to poor data quality, rather than meaningful differences in our regions of interest.

For our primary seed-based analyses of interest, we first wanted to investigate whether this PMAT network did indeed exist in our developmental sample by assessing whether there was greater connectivity within the regions of these PM and AT subnetworks than connectivity between the regions in these networks. When running a two-way ANOVA assessing within vs. between connectivity and age we found a significant interaction between age and within/between connectivity (*F*(4,150) = 2.45, *p* = 0.044). However, an overall a Tukey’s HSD test revealed that there was significantly more connectivity within the regions of the PM and AT subnetworks than between the regions of these subnetworks for all age groups: 3-4 years: (*p* = .050, CI = [-.001, 0.179]); 5-years: (*p* < .001, CI = [.008, 0.254]); 7-years: (*p* < .001, CI = [0.006, 0.270]); 8-12 years: (*p* < .001, CI = [0.009, 0.256]); Adults: (*p* = 0.050, CI = [-.001, 0.174]). This result (see Figure 2) suggests that while some age-related change may exist in the connectivity of this network, there was evidence for the PM and AT subnetworks in all age-groups.

**Figure 2.**
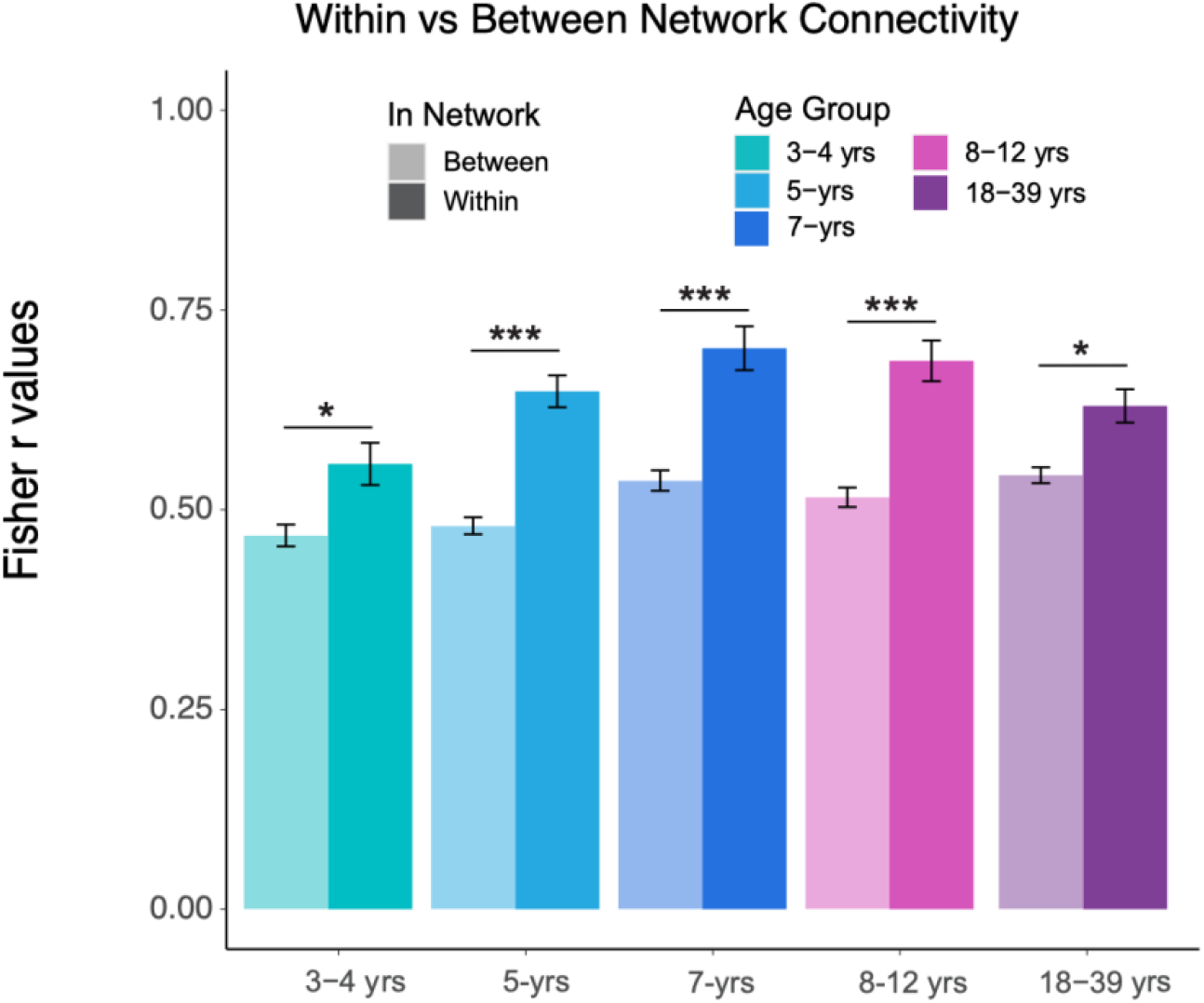
PMAT Within-Between Network Connectivity by Age Group. All five age-groups demonstrated significantly higher within than between PMAT network fisher r values. Darker colored bars represent within and lighter represent between network connectivity for the PMAT network overall. Significance markers indicate results of simple effects tests of within vs between connectivity differences (p< .05 *; p< .005 **, p <.0005 **).

Next, we wanted to look at the development of these PM and AT subnetworks separately, as well as their connectivity to the hippocampus. We ran two separate ANOVAs looking at the average connectivity of the regions *within* these subnetworks separately for PM and AT, and connectivity with the hippocampus as a function of age. For the within-network ANOVA, we found an interaction by subnetwork and age (*F*(4,150) = 4.97, *p* < .0001). Post-hoc Tukey’s HSD tests revealed that AT network connectivity did not change with age (See Figure 3A, and Supplemental Table 1 for full Tukey’s HSD results). However, there was age-related change in the PM subnetwork (Figure 3B) such that connectivity in the PM significantly increased from 3-4 to 7 years of age: (*p* = .006, CI = [0.025, 0.290]), before decreasing again between 8-12 years and adulthood: (*p* = .003, CI = [0.029, 0.26]).

**Figure 3.**
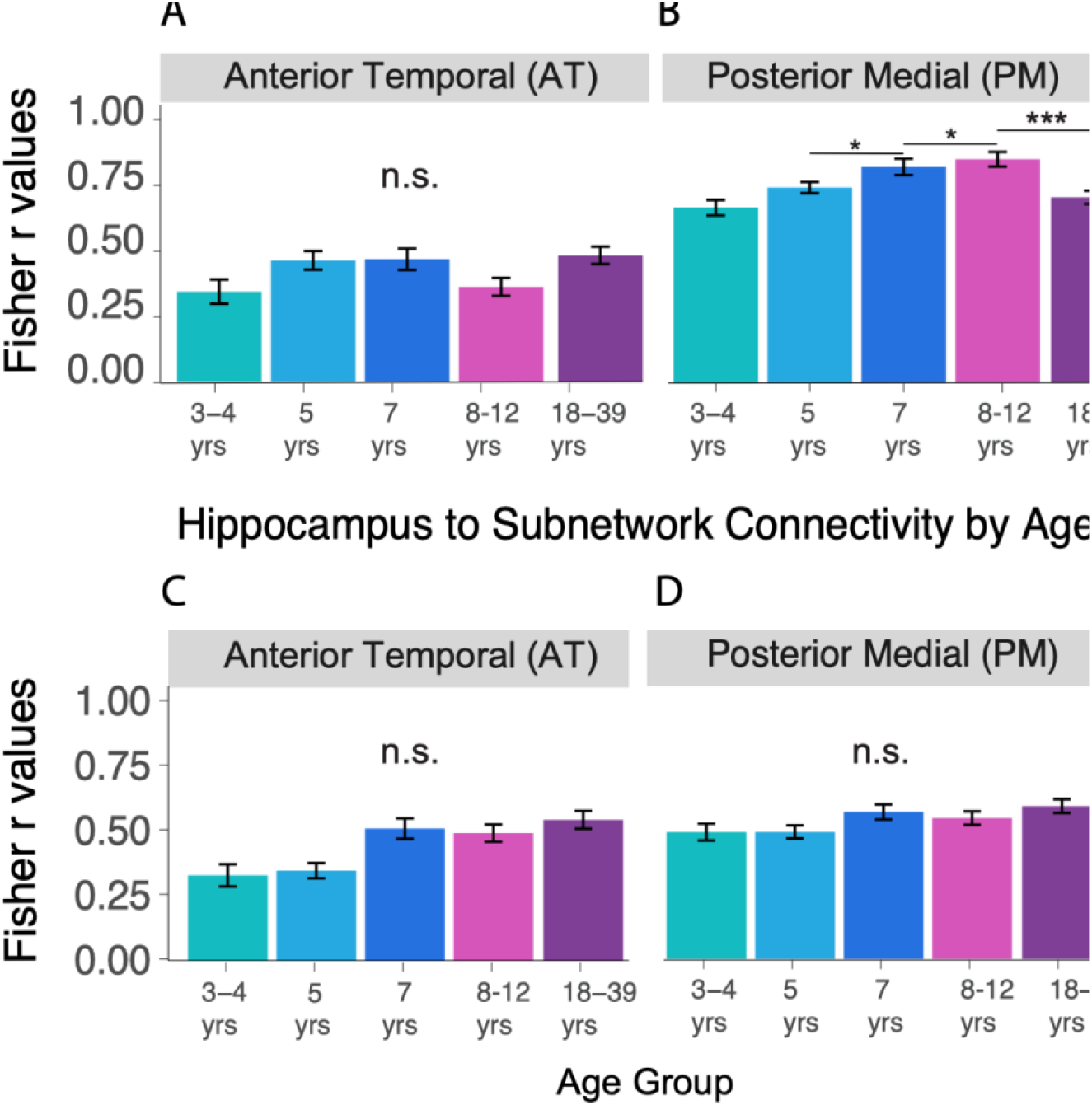
PM/AT Subnetwork and Whole Hippocampal-Subnetwork Connectivity by Age. Bars represent average fisher r correlation values for each age group by subnetwork for within subnetwork connectivity and hippocampal to subnetwork connectivity. The within-subnetwork connectivity ANOVA did demonstrate a significant interaction between sub-network and age (Panels A and B). Whereas the hippocampus-subnetwork AOVA did not yield a significant interaction between subnetwork and age. All results are plotted regardless for comparison purposes. A) Average correlations for within the AT subnetwork connectivity did not significantly change across development from Tukey’s HSD test. B). Average correlations for the within the PM subnetwork connectivity did show overall higher connectivity and with increasing correlations until 8-12 years of age from Tukey’s HSD test. C) Average correlations for the whole hippocampus to AT network. Interaction between subnetwork and age was not significant. D) Average correlations for the hippocampus to PM network. Interaction between subnetwork and age was not significant. All significance markers denote significant findings from follow-up Tukey’s HSD test and non-significant interaction ( n.s. = non-significant, *p* < .05 *; *p* <.0005 ***).

For the hippocampal-PMAT connectivity analysis we did find a significant main effect of sub-network (F(1,150) = 2.43, *p* < .001) with significantly higher PM (*M* = 0.54, *SD* = 0.36) than AT subnetwork connectivity with the hippocampus (*M* = 0.44, *SD* = 0.32). We also found a main effect of age group (*F*(4,150) = 4.08, *p* < .001) such that there was a significant difference particularly from 3-4 to 7-8 years (*p* = .003, CI = [0.029, 0.215]) as well as 5-years to 7-years of age (*p* = .005, CI = [0.022, 0.204]) after which there are no differences in age groups. However, we did not find a significant interaction (*F*(4,150) = 1.57, *p* = .179) between subnetwork and age groups (Figure 3C and 3D.)

One reason the age by subnetwork interaction for the hippocampal-PMAT ANOVA may not have been significant is that previous work shows that the long-axis of the hippocampus develops at different rates (DeMaster et al., 2014; Gogtay et al., 2006; Lee et al., 2014), subserves separate memory functions with age (Vijayarajah & Schlichting, 2024), and connects differentially with the PM and AT subnetworks across development (Xie et al., 2024). To that end, we next assessed connectivity between the anterior (head) and posterior (tail) hippocampus and the PM and AT subnetworks across development. For comparison purposes, we plot the difference between anterior and posterior hippocampal connectivity to both subnetworks (Figure 4).

**Figure 4.**
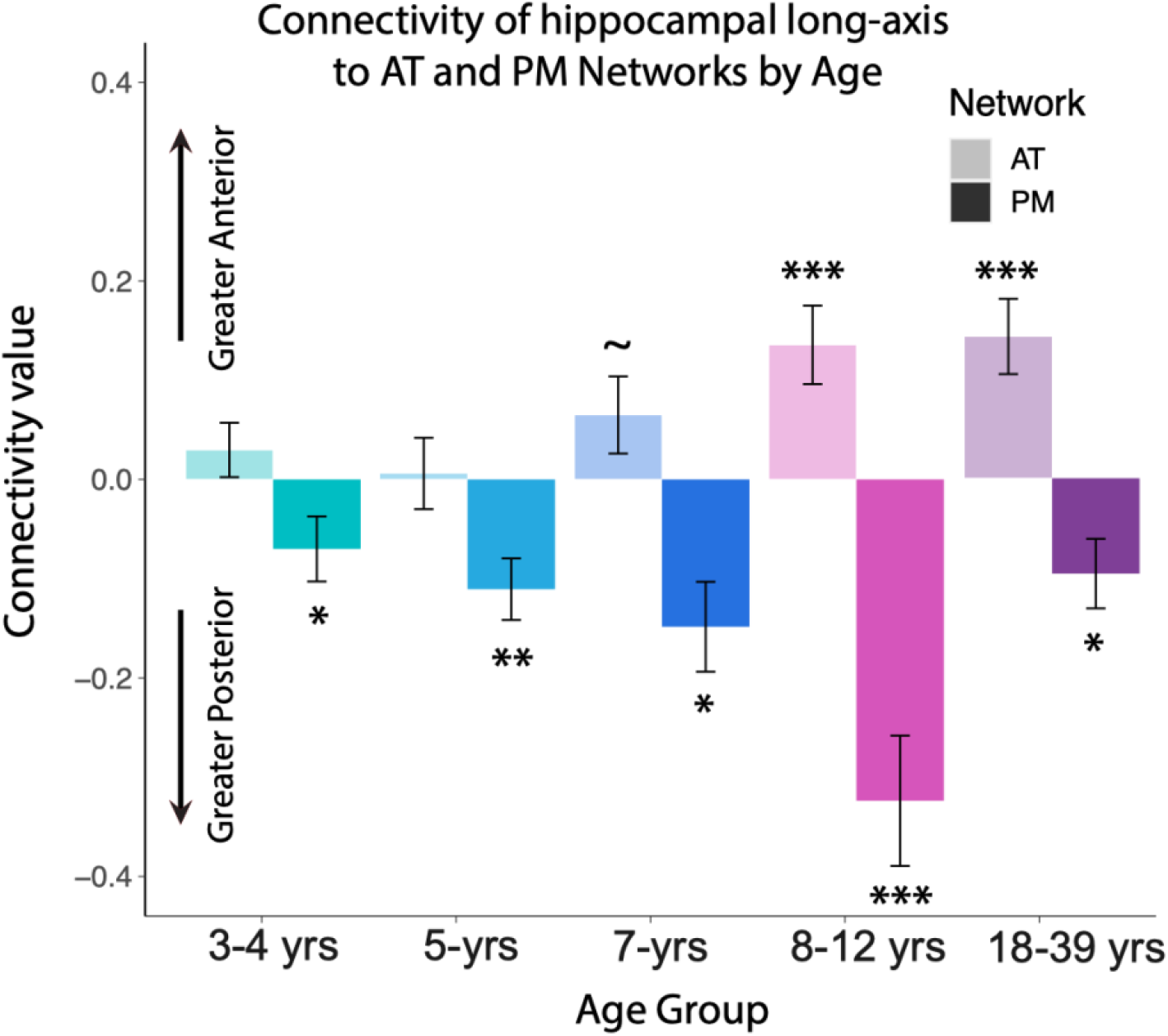
Difference in connectivity for anterior/posterior hippocampal axis and PM/AT sub networks by age. Plotted is a difference score calculated by subtracting age-group averaged anterior – posterior hippocampal fisher r correlation values to the AT and PM subnetwork respectively. Higher (more positive) values on the y-axis indicate greater anterior than posterior connectivity on average to a given network, and lower (more negative) values on the y-axis indicate greater posterior than anterior connectivity on average to a given network. Darker bars are average difference scores for the PM network by age and lighter bars are average difference scores for the AT network by age. There was significantly greater posterior connectivity (significantly negative values) to the PM network in all age groups, demonstrating PM – posterior hippocampal connectivity specificity at all ages. However, there was significantly more AT – anterior hippocampal connectivity specificity (significantly positive values) only after ages 7 years of age. All significance markers denote significant t-test differences from zero by age group ( *p* = .05 ∼, *p* < .05 *; *p* < .005 **, *p* <.0005 ***).

We ran a two-way ANOVA assessing connectivity difference scores (anterior-posterior hippocampal) by AT and PM subnetworks and age-group. We found an interaction between age and subnetwork (*F*(4,150) = 6.837, *p* < .001). Follow up Tukey’s HSD tests revealed a significant difference in (anterior – posterior) connectivity by subnetwork such that the difference in average hippocampal connectivity to the AT subnetwork was positive on average (more anterior: *M* = 0.08, *SD* = 0.20), and negative (more posterior: *M* = −0.15, *SD* = 0.26) to the PM subnetwork on average (*p*<.001, CI = [-0.280, −0.177]) (See Supplemental Table 2 for Full Tukeys HSD results). Notably, we also were interested in when the connectivity significantly differentiates between anterior and posterior hippocampus (i.e. specializes towards anterior/posterior for a sub-network).

Employing a one-way t-test against zero, we saw that the difference in anterior/posterior hippocampal connectivity for the AT subnetwork is no different than zero for the 3-4 year olds: (*t*(30) = 1.08, *p* = 0.143) or 5-year-olds: (*t*(33) = 0.17, *p* = 0.43), but is trending towards being different than zero by 7 years of age (*t*(22) = 1.67, *p* = 0.05) ––after which time the connectivity to AT remains significantly more anterior for 8-12 year olds: (*t*(33) = 3.43, *p* < 0.001) and adults: (*t*(27) = 3.77, *p* < 0.001). Meanwhile, the difference in anterior - posterior connectivity to the PM subnetwork is significantly different than zero even in all age groups –– 3-4 year olds: (*t*(30) = −2.135, *p* = 0.02); 5-year-olds: (*t*(33) = −3.570, *p* < 0.005); 7-year-olds: (*t*(22) = −3.276, *p* = 0.005); 8-12 year olds: (*t*(33) = −4.932, *p* < 0.0005); and adults: (*t*(27) = −2.74, *p* = 0.005). Overall we find that the AT network specializes its’ connectivity to the anterior hippocampus around or just after 7 years of age, whereas the PM network appears to be specialized in its connectivity to posterior hippocampus from 3-4 years onward, despite some developmental change in the overall level of connectivity.

## DISCUSSION

This study sought to characterize the stability of hippocampal responses across age and how changes in hippocampal synchrony coincided with connectivity changes between the hippocampus and cortical memory systems. We found that hippocampal synchrony is relatively stable across development except in 7-year-olds –– which was the only age group that failed to show inter-group synchrony in their hippocampal responses. These hippocampal responses instead bifurcated: about half of the 7-year-old group had a less or more mature hippocampal response, and only one 7-year-old participant had a hippocampal response most similar to their own age-group. We also found that an Anterior Temporal (AT) cortical network that supports semantic memory is early maturing, but only around 7 years of age does this AT network start to interact with specifically with the anterior hippocampus as it does in adults. Taken together, these results suggest that middle childhood may be an inflection point for hippocampal reorganization: whereby cortical brain networks that may support semantic memory behaviors are early maturing, but do not integrate with a changing hippocampus until middle childhood.

These findings build upon and extend prior work showing protracted hippocampal development patterns. Prior fMRI work shows that hippocampal systems are relatively late to mature (Ghetti & Fandakova, 2020; Ghetti & Lee, 2011), as are hippocampal interactions with lateral prefrontal cortices (Riggins et al., 2020). However, prior work focused on task-based responses to static memoranda, which precluded the ability to look at more ecologically-valid responses to less-constrained stimuli. Moreover, prior work focused mostly on episodic memory systems by only querying hippocampal activation or hippocampal connectivity within a relatively narrow range of ROIs representing sensory regions and/or task-relevant PFC responses (see Fandakova et al., 2017). While a recent study looked at connectivity between the hippocampus and broader cortical memory networks, they only modeled developmental change linearly in an older age-range; potentially missing non-linear idiosyncrasies of responses between age-groups (Xie et al 2024). Here, by leveraging ISC approaches in conjunction with functional connectivity with broad cortical memory networks, we were able to isolate an inflection point at 7 years of age, wherein there are dramatic shifts in the uniformity of hippocampal responses that coincide with connectivity changes between the hippocampus and large-scale memory networks.

Although memory was not explicitly tested in this study, our findings speak to theories about the development of neural systems underlying episodic and semantic memory systems. Prominent adult memory theories propose that episodes precede and help structure semantic knowledge (Kumaran et al., 2016; McClelland et al., 1995), but how episodic and semantic memory systems develop is still being explored. While some recognition memory behaviors appear as early as the first years of life (Bauer, 2002; Fagan, 1974) detailed episodic recall does not match adults until at least middle childhood (Ghetti & Lee, 2011; Riggins et al., 2020). In contrast, some studies demonstrate that semantic memory appears relatively mature early in life: children as young as 4 years demonstrate adult-like semantic organization and concept-based reasoning (Keresztes et al., 2018; Ngo et al., 2021) and can extract generalizations across experiences even when episodic memory is poor (Bauer et al., 2016; Varga et al., 2016). Indeed, these robust early semantic behaviors may be scaffolded on imprecise episodic systems which provide enough information to extracting generalizations and gist-representations (Sloutsky et al., 2025). However, these developmental relationships amongst episodic and semantic memory behaviors are not always so straightforward; some evidence suggests young children sometimes rely on more precise episodic details and thus generate fewer false-memories compared to adults who are more likely to respond to semantic-lures and thus false alarm more frequently (Brainerd et al., 2008). In total, behavioral evidence is quite mixed on how and when the memory functions from the episodic and semantic memory systems scaffold the other across development.

We provide evidence that the brain networks hypothesized to underlie semantic memory matures earlier than its interactions with a key region (the hippocampus) important for episodic memory encoding (Davachi, 2006; Kim, 2011) as well as translation between episodes and semantics (Kumaran et al., 2016; McClelland et al., 1995). Using a more systems neuroscience-based approach, we query well-defined networks as opposed to behavior. Our results show that the cortical brain network thought to support semantics reaches maturity early but does not synchronize with the hippocampus until middle childhood. Interestingly, these findings dovetail with other neuroscience-based inquiries done in rodent models, showing that in early childhood the hippocampus may be more oriented towards overgeneralization, which could be a neural manifestation of semantic-based memories (Elliott & Richardson, 2019).

Our findings also help contextualize the development episodic memory specifically across childhood. Prior developmental theories have postulated that middle childhood may be the offset of infantile amnesia –– the phenomenon where as adults we recall only a very sparse number of autobiographical childhood memories (Bauer & Leventon, 2013; Josselyn & Frankland, 2012). Interpreted through our current findings, shared information processing across the hippocampus and semantic memory networks may underlie adult-like episodic memories. This coincidence in timescales is actually quite surprising as the current dataset looked at brain responses during free movie watching and did not query memory retrieval explicitly. In this way, our findings lend support to previous models of memory development (Bauer, 2015), such that children may not be able to encode adult-like memories due to constraints of integrating semantic information with incoming sensory information entering the hippocampus.

This interpretation of our data supports the idea that while semantic brain networks exist early, translation between episodes and semantics may be delayed due to hippocampal reorganization. Some form of episodic and semantic memory may be able to exist independently, but are not yet communicating with one another via the hippocampus.

This has implications for bi-directional theories of memory interaction (Renoult et al., 2019): where semantic systems may initially develop independently, but later become interdependent with episodic systems. This leads to testable behavioral predictions: early in development children may be predisposed to the integration of information across episodes and may easily form gist-representations. Meanwhile, while some sporadic details of episodes may be available, they are constrained to an independent episodic system. This may lead to more episodic forgetting overall as there is little semantic context available to scaffold episodic details until these two systems learn to integrate and share information. These predictions may be worthy of future research integrating behavioral and neuroimaging techniques across wide age ranges.

While the results presented here offers a framework for how and when hippocampal and cortical memory systems begin to integrate across development, there are some notable limitations to the dataset and our approach. Firstly, the dataset used contains no behavioral measurements of memory, so hypotheses about whether hippocampal integration with cortical semantic memory systems directly relates to episodic and semantic behaviors are speculative and yet to be tested. Likewise, we cannot be sure that the changes seen here do not underlie memory behaviors (like pattern completion) that are shared by multiple memory systems. A future avenue for research would be to tie these methods looking at changes in hippocampal stability and connectivity directly to behavioral memory tasks that disambiguate between these multiple functions. Furthermore, the maturity differences we see in 7-year-olds hippocampal responses with the ISC measurement could be due to individual differences not measured in this dataset. Early life stress for example has been shown to influence temporarily impact the speed of hippocampal maturity (Humphreys et al., 2019; Tottenham, 2009), but this was outside the scope of this investigation.

Secondly, the dataset is cross-sectional which limits our ability to ask, for example, whether 7 year olds, who are early or late maturing in their hippocampal responses, remain early or late maturing for their age group, and whether this is predictive of future memory behaviors. Likewise, while we endeavored to select a dataset with a relatively wide age-range, this dataset has notable gaps: there are no six year olds included, so whether the changes reported occur precisely around 7-years, or start In 6-year olds remains is unknown. Likely, this is a gradual change that happens around 7-years rather than punctate –– indeed, when the main findings from this dataset are plotted continuously, 7-8 year olds look fairly similar before diverging nearing age 9 (see Supplemental Figure 1 and 2). Furthermore, there are no adolescents in this dataset (though see Xie et al., 2024 for an adolescent age range) where there has been evidence that hippocampal changes are related to unique memory-based behaviors in this age range (Davidow et al., 2016). It would be interesting to zoom in on adolescence to interrogate whether changes in hippocampal synchrony and their connectivity with broader cortical memory networks are related to increases in episodic (Ghetti & Angelini, 2008) or semantic (Pudhiyidath et al., 2020) or other long-term memory behaviors (Skalaban et al., 2022) that show non-linear changes from adolescence into adulthood.

Overall, this study harnessed inter-subject correlation methods, extending prior work on the development of the hippocampus, to pinpoint a potential functional inflection point at 7 years of age. This age-range is also when connectivity between the hippocampus and an anterior temporal cortical memory network began to look more adult-like, despite the anterior temporal memory network showing mature connectivity even in toddlers. These findings may call into question whether semantic memory actually precedes the development of episodic memory functions and suggests that the two systems may operate independently early in development. Instead, neural integration of the two systems in the brain may not begin until middle childhood and may be a potential source of more adult-like memory behaviors.

## MATERIALS AND METHODS

### Dataset

To test our hypotheses, we utilized a publicly available dataset including functional MRI data collected in 3-12 year old children and adults (Richardson et al., 2018; OpenNeuro) (N = 155). This data was obtained from the OpenNeuro database. Its accession number is ds000228. All subjects watched Disney Pixar’s “Partly Cloudy” a short 5 minute long film in the scanner sans any other task. The total length of the initial experiment was 5.6 minutes TR = 2s IPS = 168 TRs, and the film began after 10s of rest followed by 10s of opening credits. The publicly available version of the dataset accessed here excluded five child subjects for excessive head motion – which the original authors defined as having greater than one-third of their time points for which there was either >2mm composite motion relative to the previous timepoint or a fluctuation in global signal that exceeded a threshold of three s.d. from the global mean signal. A full description of the dataset and experimental parameters can be found in the original paper (Richardson et al., 2018).

### Sample

The preprocessed dataset included in this paper contained 150 subjects which we split up into five age groups (N = 31, age mean = 4.06 yrs, age range = 3.5 – 4.86 yrs), 5-year-olds (N = 34, age mean = 5.51 yrs, age range = 5.01 – 5.99 yrs), 7-year-olds (N = 23, age mean = 7.54, age range = 7.00 – 7.96 yrs), 8-12 year olds (N = 34, age mean = 9.77, age range = 8.02 – 12.30 yrs) and adults (N = 28, age mean = 24.78, age range = 18-39 years). It should be noted that there were no six-year olds (6.0 – 6.9 years) included in this dataset, and that the 8-12 year-old group of children includes a broader range of ages than the other groups.

We split these age groups in this way to try and accomplish two goals: Firstly, to represent key periods of change in memory development from toddlers to school-age children (Bauer et al., 2016; Riggins et al., 2016), and from early childhood to middle childhood (Canada et al., 2020; Ghetti & Lee, 2011; Riggins et al., 2020). Secondly, we endeavored to split the sample into as close of equal age-bins as possible. This could have been accomplished in a myriad of ways, but as the number of participants who were 5-years of age exceeded any the number of participants in any other year (n = 34), we needed the closest adjacent age-groups (3-4 and 7-year-olds) to be separate from the 5-year olds so as not to overweight that group. We could have included 8-year olds in the 7-year age group: (n = 35) and let the oldest age-group include only 9-12 year olds: (n = 22) and the number of subjects in these two groups would have essentially switched. Unavoidably, one group in this sample must have a somewhat smaller number of subjects than the others, or otherwise the ages would have to be split in between years of age (with a boundary at say 8.5) which would seem fairly arbitrary and atheoretical.

However, based on observations from a continuous analysis, adding the 8-year olds to that group would not appear to change the directionality/patterns of results for the ISC (see Supplemental Figure 1) or the connectivity results (Supplemental Figure 2) one way or the other except to strengthen the current results. Therefore, we decided to complete analyses only with the age-groups we selected initially and not to re-run analyses including the 8-year olds with 7-year olds as to avoid unnecessary multiple comparisons.

### Image Preprocessing

We performed preprocessing on the BIDS formatted publicly available data using fMRIprep (Esteban et al., 2019). Results included in this manuscript come from preprocessing performed using FMRIPREP version 20.2.0 [1, 2, RRID:SCR_016216], a Nipype [3, 4, RRID:SCR_002502] based tool. Each T1w (T1-weighted) volume was corrected for INU (intensity non-uniformity) using [N4Bias Field Correction] v2.1.0 [5] and skull-stripped using antsBrainExtraction sh v2.1.0 (using the OASIS template). Brain surfaces were reconstructed using recon-all from FreeSurfer v6.0.1 [6, RRID:SCR_001847], and the brain mask estimated previously was refined with a custom variation of (Daugherty et al., 2016; Ghetti & Bunge, 2012; Gómez & Edgin, 2016; Lee et al., 2014) of Mindboggle [21, RRID:SCR_002438]. Spatial normalization to the ICBM 152 Nonlinear Asymmetrical template version 2009c [7, RRID:SCR_008796] was performed through nonlinear registration with the antsRegistration tool of ANTs v2.1.0 [8, RRID:SCR_004757], using brain-extracted versions of both T1w volume and template. Brain tissue segmentation of cerebrospinal fluid (CSF), white-matter (WM) and gray-matter (GM) was performed on the brain-extracted T1w using [17] (FSL v5.0.9, RRID:SCR_002823).

Functional data was slice time corrected using 3dTshift from AFNI v16.2.07 [11, RRID:SCR_005927] and motion corrected using mcflirt (FSL v5.0.9 [9]). This was followed by co-registration to the corresponding T1w using boundary-based registration [16] with six degrees of freedom, using bbregister (FreeSurfer v6.0.1). Motion correcting transformations, BOLD-to-T1w transformation and T1w-to-template (MNI) warp were concatenated and applied in a single step using antsApplyTransforms (ANTs v2.1.0) using Lanczos interpolation.

Physiological noise regressors were extracted applying CompCor [18]. Principal components were estimated for the two CompCor variants: temporal (tCompCor) and anatomical (aCompCor). A mask to exclude signal with cortical origin was obtained by eroding the brain mask, ensuring it only contained subcortical structures. Six tCompCor components were then calculated including only the top 5% variable voxels within that subcortical mask. For aCompCor, six components were calculated within the intersection of the subcortical mask and the union of CSF and WM masks calculated in T1w space, after their projection to the native space of each functional run. Frame-wise displacement [19] was calculated for each functional run using the implementation of Nipype.

Many internal operations of FMRIPREP use Nilearn [22, RRID:SCR_001362], principally within the BOLD-processing workflow. For more details of the pipeline: see https://fmriprep.readthedocs.io/en/20.2.0/workflows.html

### Defining Regions of Interest/Networks

We opted to align all subjects’ functional data to the same adult-defined standard space in Freesurfer and extracted a set of anatomically defined bilateral ROIs using the Desikan-Killiany Atlas (Desikan et al., 2006). For all brain regions, we used bilateral left and right ROIs and combined them into a singular non-contiguous ROI. In the ISC analyses (Figures 1) and initial connectivity analyses (Figure 3), we used the whole hippocampus and for the post-hoc connectivity analyses only (Figure 4) we divided the hippocampus into three segment along the long-axis and used the most the most anterior (head) and posterior (tail) segments (Murty et al., 2017). For cortical memory networks, we selected ROIs wherein we selected anatomical nodes that were the most aligned with functional networks defined in a recent paper looking at these Posterior Medial and Anterior Temporal networks (Barnnett et al., 2021). These included the following regions: Posterior Medial: posterior cingulate cortex, isthmus cingulate cortex, superior frontal gyrus, supramarginal gyrus; Anterior Temporal: temporopolar, lateral orbitofrontal, superior parietal).

### Inter-subject Functional Correlation Analyses

All post-preprocessing analyses were conducted in Python-3 using custom written scripts based on Nipy, nilearn and connectivity tools publicly available through the Brain Imaging Analysis Kit (BrainIAK; https://brainiak.org). First, each subjects’ time course, or vector of activity amplitudes across TRs averaged over voxels in a given ROI, was extracted. Then, in order to determine how much an individual’s hippocampal timecourse matches their given age-group, we conducted a Pearson correlation between a given subject and the average response for a given age group leaving that one subject out of the average at a time. Next, to determine how much each subject looked like other age-groups, we conducted a Pearson correlation between a given subjects hippocampal timecourse and the average timecourse of all subjects in each of the five age groups.

To evaluate whether this was fewer subjects than would be expected in their own age group than by chance, we conducted a permutation analysis assuming that each 7 year old was equally as likely to be correlated with any of the other five age groups. As some expected counts were small, we estimated p-values using Monte-Carlo simulation. To do this, we simulated labels for 23 subjects, each time randomly assigning a “winning” age-group label and then re-labeled according to whether this age group was younger, older or the same as the 7 year old group and then computed a chi-square statistic comparing simulated counts to their expected values. This was repeated 100,000 times to generate an empirical null-distribution of the test statistic. This observed chi-square statistic was compared to this null distribution to obtain a Monte Carlo p-value. This approach accounts for the unequal number of templates contributing to the younger, same-age, and older categories, but preserves the original winner-take all analysis structure.

### Seed-based Connectivity Analyses

Given our networks of interest, we calculated pairwise Pearson correlations of the timecourses between two ROIs within each network for each subject. Notably, for some analyses, this included exclusively cortical ROIs (within network only) and in some cases included both hippocampal and cortical ROIs (hippocampus/anterior/posterior hippocampus to each region in the PM or AT network).

### Group-level statistical analyses

To assess all age-related differences of our various measures of interest, we conducted two-way analyses of variance (ANOVA) using the AOV function in the stats package of R version (1.2.5042). We assessed fisher r corrected correlation values by age-group and network. For any significant ANOVA interactions, we conducted post-hoc analyses using Tukey’s HSD tests to account for multiple comparisons. In the case of the anterior/posterior hippocampal to PMAT subnetwork analyses, we first calculated average connectivity between the regions of a given subnetwork to anterior and posterior hippocampus respectively, and they subtracted the resultant fisher r-value for posterior hippocampus from the r-value for anterior hippocampus. In addition to an ANOVA looking at the interaction between subnetwork and age-group, we also then conducted a one-way t-test against zero to see if this difference score was significantly different than zero for each age-group. All significance was set to level of *p* < .05.

## Supporting information

Supplemental Analyses

