## Supplemental Analyses for "The hippocampus and cortical memory networks have an inflection point in middle childhood"

| | diff | lwr | upr | p adj |
|---|---|---|---|---|
| 3-4 year olds:AT-18-39 year olds:AT | -0.138 | -0.306 | 0.029 | 0.209 |
| 5 year olds:AT-18-39 year olds:AT | -0.019 | -0.183 | 0.144 | 1 |
| 7 year olds:AT-18-39 year olds:AT | -0.015 | -0.197 | 0.167 | 1 |
| 8-12 year olds:AT-18-39 year olds:AT | -0.12 | -0.284 | 0.043 | 0.37 |
| 18-39 year olds:PM-18-39 year olds:AT | 0.224 | 0.081 | 0.367 | 0 |
| 3-4 year olds:PM-18-39 year olds:AT | 0.184 | 0.04 | 0.329 | 0.002 |
| 5 year olds:PM-18-39 year olds:AT | 0.261 | 0.119 | 0.403 | 0 |
| 7 year olds:PM-18-39 year olds:AT | 0.34 | 0.187 | 0.493 | 0 |
| 8-12 year olds:PM-18-39 year olds:AT | 0.369 | 0.227 | 0.511 | 0 |
| 5 year olds:AT-3-4 year olds:AT | 0.119 | -0.047 | 0.285 | 0.409 |
| 7 year olds:AT-3-4 year olds:AT | 0.123 | -0.061 | 0.308 | 0.511 |
| 8-12 year olds:AT-3-4 year olds:AT | 0.018 | -0.148 | 0.184 | 1 |
| 18-39 year olds:PM-3-4 year olds:AT | 0.362 | 0.217 | 0.508 | 0 |
| 3-4 year olds:PM-3-4 year olds:AT | 0.323 | 0.176 | 0.47 | 0 |
| 5 year olds:PM-3-4 year olds:AT | 0.4 | 0.255 | 0.545 | 0 |
| 7 year olds:PM-3-4 year olds:AT | 0.478 | 0.323 | 0.634 | 0 |
| 8-12 year olds:PM-3-4 year olds:AT | 0.507 | 0.362 | 0.652 | 0 |
| 7 year olds:AT-5 year olds:AT | 0.004 | -0.176 | 0.185 | 1 |
| 8-12 year olds:AT-5 year olds:AT | -0.101 | -0.263 | 0.061 | 0.62 |
| 18-39 year olds:PM-5 year olds:AT | 0.243 | 0.102 | 0.385 | 0 |
| 3-4 year olds:PM-5 year olds:AT | 0.204 | 0.061 | 0.347 | 0 |
| 5 year olds:PM-5 year olds:AT | 0.281 | 0.14 | 0.421 | 0 |
| 7 year olds:PM-5 year olds:AT | 0.359 | 0.208 | 0.511 | 0 |
| 8-12 year olds:PM-5 year olds:AT | 0.388 | 0.248 | 0.529 | 0 |
| 8-12 year olds:AT-7 year olds:AT | -0.105 | -0.286 | 0.075 | 0.705 |
| 18-39 year olds:PM-7 year olds:AT | 0.239 | 0.077 | 0.401 | 0 |
| 3-4 year olds:PM-7 year olds:AT | 0.199 | 0.036 | 0.363 | 0.004 |
| 5 year olds:PM-7 year olds:AT | 0.276 | 0.115 | 0.438 | 0 |
| 7 year olds:PM-7 year olds:AT | 0.355 | 0.184 | 0.526 | 0 |
| 8-12 year olds:PM-7 year olds:AT | 0.384 | 0.223 | 0.545 | 0 |
| 18-39 year olds:PM-8-12 year olds:AT | 0.344 | 0.203 | 0.486 | 0 |
| 3-4 year olds:PM-8-12 year olds:AT | 0.305 | 0.162 | 0.448 | 0 |
| 5 year olds:PM-8-12 year olds:AT | 0.381 | 0.241 | 0.522 | 0 |
| 7 year olds:PM-8-12 year olds:AT | 0.46 | 0.309 | 0.612 | 0 |
| 8-12 year olds:PM-8-12 year olds:AT | 0.489 | 0.349 | 0.63 | 0 |
| 3-4 year olds:PM-18-39 year olds:PM | -0.04 | -0.158 | 0.079 | 0.988 |
| 5 year olds:PM-18-39 year olds:PM | 0.037 | -0.079 | 0.153 | 0.991 |
| 7 year olds:PM-18-39 year olds:PM | 0.116 | -0.013 | 0.244 | 0.12 |
| 8-12 year olds:PM-18-39 year olds:PM | 0.145 | 0.029 | 0.26 | 0.003 |
| 5 year olds:PM-3-4 year olds:PM | 0.077 | -0.041 | 0.194 | 0.55 |
| 7 year olds:PM-3-4 year olds:PM | 0.155 | 0.025 | 0.286 | 0.006 |
| 8-12 year olds:PM-3-4 year olds:PM | 0.184 | 0.067 | 0.302 | 0 |
| 7 year olds:PM-5 year olds:PM | 0.079 | -0.049 | 0.206 | 0.633 |
| 8-12 year olds:PM-5 year olds:PM | 0.108 | -0.007 | 0.223 | 0.087 |
| 8-12 year olds:PM-7 year olds:PM | 0.029 | -0.099 | 0.157 | 0.999 |

**Supplemental Table 1** Full Tukey's HSD result for interaction in ANOVA looking at Connectivity Within in the PMAT network by subnetwork (PM/AT) and Age Group.

| | diff | lwr | upr | p adj |
|---|---|---|---|---|
| 3-4 year olds:AT-18-39 year olds:AT | -0.113 | -0.3 | 0.074 | 0.647 |
| 5 year olds:AT-18-39 year olds:AT | -0.137 | -0.32 | 0.046 | 0.337 |
| 7 year olds:AT-18-39 year olds:AT | -0.078 | -0.28 | 0.124 | 0.966 |
| 8-12 year olds:AT-18-39 year olds:AT | -0.007 | -0.19 | 0.175 | 1 |
| 18-39 year olds:PM-18-39 year olds:AT | -0.239 | -0.43 | -0.048 | 0.003 |
| 3-4 year olds:PM-18-39 year olds:AT | -0.213 | -0.4 | -0.026 | 0.012 |
| 5 year olds:PM-18-39 year olds:AT | -0.253 | -0.436 | -0.071 | 0.001 |
| 7 year olds:PM-18-39 year olds:AT | -0.291 | -0.493 | -0.09 | 0 |
| 8-12 year olds:PM-18-39 year olds:AT | -0.467 | -0.649 | -0.284 | 0 |
| 5 year olds:AT-3-4 year olds:AT | -0.024 | -0.202 | 0.154 | 1 |
| 7 year olds:AT-3-4 year olds:AT | 0.035 | -0.162 | 0.232 | 1 |
| 8-12 year olds:AT-3-4 year olds:AT | 0.106 | -0.072 | 0.284 | 0.671 |
| 18-39 year olds:PM-3-4 year olds:AT | -0.126 | -0.313 | 0.061 | 0.495 |
| 3-4 year olds:PM-3-4 year olds:AT | -0.1 | -0.282 | 0.082 | 0.767 |
| 5 year olds:PM-3-4 year olds:AT | -0.14 | -0.318 | 0.038 | 0.265 |
| 7 year olds:PM-3-4 year olds:AT | -0.178 | -0.375 | 0.019 | 0.116 |
| 8-12 year olds:PM-3-4 year olds:AT | -0.353 | -0.531 | -0.176 | 0 |
| 7 year olds:AT-5 year olds:AT | 0.059 | -0.134 | 0.252 | 0.994 |
| 8-12 year olds:AT-5 year olds:AT | 0.13 | -0.044 | 0.303 | 0.343 |
| 18-39 year olds:PM-5 year olds:AT | -0.102 | -0.285 | 0.081 | 0.747 |
| 3-4 year olds:PM-5 year olds:AT | -0.076 | -0.254 | 0.102 | 0.937 |
| 5 year olds:PM-5 year olds:AT | -0.117 | -0.29 | 0.057 | 0.501 |
| 7 year olds:PM-5 year olds:AT | -0.154 | -0.348 | 0.039 | 0.248 |
| 8-12 year olds:PM-5 year olds:AT | -0.33 | -0.503 | -0.156 | 0 |
| 8-12 year olds:AT-7 year olds:AT | 0.071 | -0.123 | 0.264 | 0.977 |
| 18-39 year olds:PM-7 year olds:AT | -0.161 | -0.363 | 0.041 | 0.248 |
| 3-4 year olds:PM-7 year olds:AT | -0.135 | -0.332 | 0.062 | 0.471 |
| 5 year olds:PM-7 year olds:AT | -0.175 | -0.369 | 0.018 | 0.112 |
| 7 year olds:PM-7 year olds:AT | -0.213 | -0.425 | -0.002 | 0.045 |
| 8-12 year olds:PM-7 year olds:AT | -0.389 | -0.582 | -0.195 | 0 |
| 18-39 year olds:PM-8-12 year olds:AT | -0.232 | -0.414 | -0.049 | 0.003 |
| 3-4 year olds:PM-8-12 year olds:AT | -0.206 | -0.383 | -0.028 | 0.01 |
| 5 year olds:PM-8-12 year olds:AT | -0.246 | -0.42 | -0.072 | 0 |
| 7 year olds:PM-8-12 year olds:AT | -0.284 | -0.477 | -0.091 | 0 |
| 8-12 year olds:PM-8-12 year olds:AT | -0.459 | -0.633 | -0.286 | 0 |
| 3-4 year olds:PM-18-39 year olds:PM | 0.026 | -0.161 | 0.213 | 1 |
| 5 year olds:PM-18-39 year olds:PM | -0.015 | -0.197 | 0.168 | 1 |
| 7 year olds:PM-18-39 year olds:PM | -0.052 | -0.254 | 0.149 | 0.998 |
| 8-12 year olds:PM-18-39 year olds:PM | -0.228 | -0.41 | -0.045 | 0.004 |
| 5 year olds:PM-3-4 year olds:PM | -0.041 | -0.218 | 0.137 | 0.999 |
| 7 year olds:PM-3-4 year olds:PM | -0.078 | -0.275 | 0.119 | 0.96 |
| 8-12 year olds:PM-3-4 year olds:PM | -0.254 | -0.432 | -0.076 | 0 |
| 7 year olds:PM-5 year olds:PM | -0.038 | -0.231 | 0.156 | 1 |
| 8-12 year olds:PM-5 year olds:PM | -0.213 | -0.387 | -0.039 | 0.004 |
| 8-12 year olds:PM-7 year olds:PM | -0.175 | -0.369 | 0.018 | 0.113 |

**Supplemental Table 2** Full Tukey's HSD result for interaction in ANOVA looking at differences in connectivity in Anterior and Posterior Hippocampus by subnetwork (PM/AT) and Age Group.

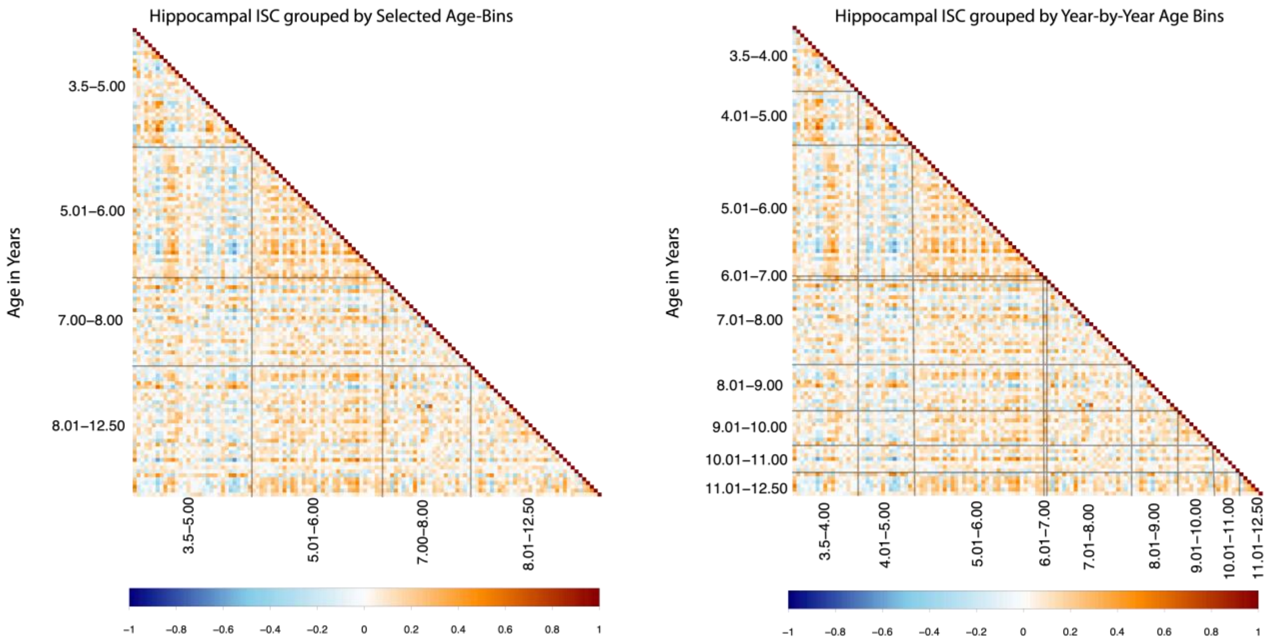


**Supplemental Figure 1: ISFC matrices between all developmental subjects plotted continuously with groupings by selected age-bins and year-to-year bins.**

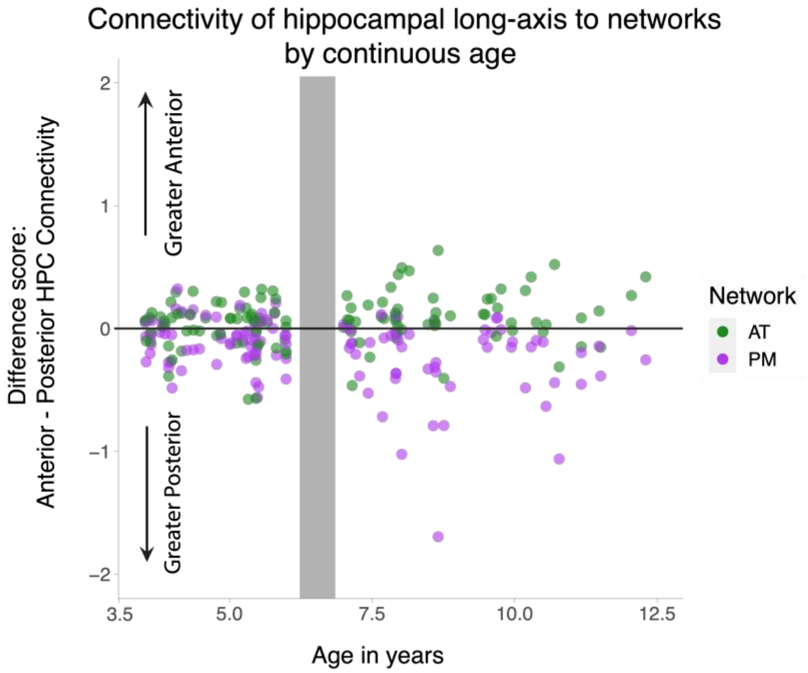


**Supplemental Figure 2: Connectivity of the hippocampal long axis to PM and AT networks plotted continuously by age.**
